# IgE-producing cells on the move: CCR2 is a key regulator of IgE^+^ plasma cell migration

**DOI:** 10.64898/2025.12.18.695109

**Authors:** Zihan Liu, Pavel Tolar, Faruk Ramadani

## Abstract

**Background:** Immunoglobulin E (IgE) plays a fundamental role in the pathogenesis of allergic disease, including asthma. The IgE-producing plasma cells (PCs) are thought to persist indefinitely, providing a sustained source of allergen-specific IgE. Although these cells can accumulate in the bone marrow (BM), after prolonged allergen exposure, their frequency remains remarkably low, and the mechanisms that regulate their migration are poorly understood.

**Objective:** To investigate the chemokine receptor profile and the migration potential of the human IgE-producing cells.

**Methods:** Tonsil B cells were stimulated with IL-4 and anti-CD40 to induce class switching to IgE and IgG1. The chemokine receptor profile of IgE^+^ and IgG1^+^ switched cells was determined using flow cytometry and migration towards relevant chemokines was quantified using transwell chemotaxis assays. Chemokine expression was also validated by re-analysis of a published single cell RNA sequencing (scRNAseq) dataset of PCs isolated from nasal polyps (NP) of patients with allergic fungal rhinosinusitis.

**Results:** IgE_⁺_ PCs exhibit significantly reduced expression of the BM-homing chemokine receptor CXCR4 and impaired migration towards its ligand, CXCL12. While IgE^+^ PCs can upregulate CCR10 and respond to its ligand, CCL28, this behaviour is similar to IgG1^+^ PCs. Strikingly, however, IgE_⁺_ PCs selectively upregulate CCR2 and migrate robustly towards its ligand CCL2. Re-analysis of NP scRNAseq data confirmed that IgE_⁺_ PCs express significantly higher levels of CCR2 compared with PCs of all other isotypes.

**Conclusions:** These findings identify CCR2 as a key regulator of IgE_⁺_ PC migration and provide insights into their homing preferences that may shape the nature of the IgE responses.

## Introduction

Allergen-specific IgE antibodies are central to the allergic response (1,2). They bind to the high affinity IgE receptor (FcεRI) on mast cells and basophils, which when bound by allergens trigger these effector cells to release inflammatory mediators responsible for the immediate hypersensitivity reaction (1). IgE antibodies are produced by plasma cells (PCs) after immunoglobulin class switching to IgE in B cells of other Ig isotypes. The location and longevity of these IgE-producing PCs have long been a contentious point (3–7).

The migration and tissue localisation of B cells and PCs are primarily dictated by chemokines and their receptors (8). The expression of chemokine receptors changes during B cell development and PC differentiation, and certain subsets of B cells and PCs express specific chemokine receptors that reflect their specialised functions(8,9). For example, CXCR5 and CXCR4 play an important role in shaping the B cell dynamics during the germinal centre (GC) response and their positive selection during PC differentiation (10,11), whereas CCR10 and CCR9, which are expressed at high levels on IgA^+^ PCs, are essential for migration and localisation of these cell to mucosal tissues (12–14). Similarly, the reduced expression of CXCR4, CXCR5, CCR7, CXCR3 and CCR6 on IgG4^+^ B cells has been proposed to reduce their presence in GCs of secondary lymphoid organs, contributing to the shorter lifespan of IgG4^+^ PCs compared to IgG1^+^ PCs (15).

IgE^+^ PC’s have also been reported to have reduced migratory ability, and this has been suggested to influence the dynamics of the IgE antibody response (3,4,6). Compared to IgG1^+^ PCs, mouse IgE^+^ PCs respond poorly to CXCL12 (6), a chemokine that regulates migration and retention of PCs into the BM survival niches (16,17). Indeed, only a few IgE^+^ PCs can be detected in the BM of mice following chronic allergen exposure, and they have also been shown to accumulate in the spleen, draining lymph nodes and lungs of mice, where they provide high local titres of allergen-specific IgE (5,7,18–20).

The location of PCs determines their access to survival signals and, consequently, their long-lived potential (21). In addition to the BM (5,7,18), both the lung (7) and the spleen (5) were also suggested to harbour long-lived IgE^+^ PCs. More recently, two mouse time stamping studies demonstrated that the vast majority of long-lived IgE^+^ PCs, generated during chronic allergen exposure, are found outside the BM (19,20). Due to limited access to human tissue, less is known about IgE^+^ PC localisation in humans. Nevertheless, human IgE^+^ PCs can be found in the gastrointestinal tract (22), and in the nasal polyps (NPs) of allergic individuals (23,24), sites where IgE^+^ PCs are produced (24,25). Furthermore, a small number of IgE^+^ PC can be detected in the BM of allergic individuals (7,18). This scarcity of long-lived BM IgE^+^ PCs may potentially be explained by their impaired migration to CXCL12, which was initially attributed to the reduced CXCR4 expression in IgE^+^ PCs compared with IgG1^+^ PCs (6). However, subsequent studies have reported conflicting findings (18,19). In mice, one study showed reduced CXCR4 on IgE^+^ PCs in the draining lymph nodes and BM (18), whereas another reported comparable CXCR4 levels to IgG1^+^ PCs (19). Only in the spleen the long-lived IgE^+^ PCs had significantly reduced CXCR4 compared with IgG1^+^ PCs (19). In humans, IgE^+^ PCs from the NPs of allergic individuals have higher CXCR4 transcript levels compared to other non-IgE^+^ PCs (23), while BM IgE^+^ PCs from allergic donors showed reduced CXCR4 transcripts (18). Together, these discrepancies indicate that factors beyond CXCR4 likely influence the distinct localisation of IgE^+^ PCs. How these cells home to and persist within certain tissues remains unclear.

To investigate the migration potential of IgE^+^ cells, here, we used a previously characterised human IgE class-switching tonsil B cell culture system and investigated their chemokine receptor profile and migration responses to relevant chemokines. We further validated the key findings by re-analysing a published single cell RNA sequencing (sc-RNAseq) dataset of PCs from human NPs(23).

## Materials and methods

### Ethics

Following full informed written consent, tonsils were obtained from patients undergoing routine tonsillectomies for recurrent tonsillitis or airway obstruction at the Guy’s and St Thomas’s NHS Fundation Trust. The study was conducted at and in accordance with the recommendations of King’s College London and Guy’s and St Thomas’s NHS Fundation Trust and the protocol was approved by the London Bridge Research Ethics Committee (REC number 08/H0804/94).

### Isolation of tonsil B cells

Human B cells were isolated from freshly surgically removed tonsils using 2-aminoethylisothiouronium bromide-treated sheep red blood cells (TCS Biosciences Ltd) as previously described (26,27). The B cells were then further enriched by magnetic cell sorting using the MojoSort Human pan B Cell Isolation Kit according to the manufacturer’s instructions (BioLegend).

### Class switching cultures

Purified tonsil B cells were then cultured at 0.5 x10^6^ cells/mL in RPMI 1640 (Lonza) containing penicillin (100 IU/mL), streptomycin (100 μg/mL) and glutamine (2 mM, Invitrogen) and 10% Foetal Calf Serum (Hyclone; Perbio Biosciences). To induce class switching to IgE, cultures were stimulated with IL-4 (400 IU/ml; R&D Europe Systems Ltd) and anti-CD40 antibody (1 μg/mL; G28.5; BioXCell), Transferrin (35 µg/mL) (Sigma-Aldrich) and insulin (5 µg/mL) (Sigma-Aldrich). The cells were then incubated for up to 12 days at 37°C with 5% CO2.

### Flow cytometry

After 12 days of class-switching culture, cells were harvested and washed with FACS buffer (PBS containing 5% goat serum). The cells were then resuspended in FACS buffer and blocked for 10 mins with FcR block (Miltenyi Biotec). To determine the expression levels of different chemokine receptors the cells were surface stained, as per manufactures recommendations, with BV421 anti-human CD138 (Biolegend, MI15) and either APC anti-human CXCR4 (Biolegend, 12G5), APC anti-human CCR1(Biolegend, 5F10B29), APC anti-human CCR2 (Biolegend, K036C2), or APC anti-human CCR10 (Biolegend, 6588-5) for 15-20 mins on ice. Following washing, the cells were fixed with 2% paraformaldehyde (PFA) for 10 mins at 37°C, and then permeabilized with PBS containing 0.5% saponin and 0.5% Triton x100 for 20 mins at room temperature in the dark. The permeabilised cells were washed with FACS buffer and stained with PE/Cy7 anti-human IgE (Biolegend, MHE-18; used at 1/300) and PE anti-human IgG1 (Miltenyi Biotec, IS11-12E4.23.20; used at 1/800) for 30 mins in the dark. Cells were then washed and resuspended in 200µl FACS buffer and acquired using BD LSRFortessa. The data were analysed using FlowJo software (Tree Star Inc, USA).

### Transwell Chemotaxis assay

To investigate the migration capacity of IgE^+^ and IgG1^+^ cells in response to different chemokines we used the transwell migration assay. The assay was performed using 6.5 mm Transwell® with 5.0 µm Pore Polycarbonate Membrane Insert (Corning, USA). To the bottom chamber of the transwell we added 600μl of RPMI 1640 medium only (negative control) or 600μl of RPMI 1640 supplemented with various chemokines. More specifically, we tested the migration of IgE^+^ and IgG1^+^ cells in response to recombinant human CXCL12 (R&D systems; 300ng/mL), CCL2 (Biolegend; 10ng/mL, 100ng/mL and 300ng/mL) and CCL28 (R&D systems; 300ng/mL and 1.5ug/mL). The class switched cells from day 12 of the IL4 and anti-CD40 stimulated cultures, were harvested and washed with RPMI 1640 media by centrifugation. The pelleted cells were then resuspended at 10x10^6^ cells per mL of RPMI 1640 media. 100μl of this cell suspension was added to the top chamber (insert) of the Transwell and cells were allowed to migrate to the bottom chamber for 3 h at 37 °C, with 5% CO2.

### Quantification of cell migration

To quantify the number of migrated IgE^+^ and IgG1^+^ cells in the bottom chambers, after 3h of migration, we removed the top chambers from the plates using forceps and added 100μl of Precision Count beads (Biolegend) to the bottom chambers. After thorough mixing, the suspension from the bottom chambers were transferred to FACS tubes and surface stained for CD138 and intracellularly for IgE and IgG1 as above. The cells were then acquired using BD FACSCanto or BD LSRFortessa. To calculate the number of cells that migrated we divided the number of acquired cells by the number of counted beads and multiplied this number by 100,000 (the total number of Precision Count beads added to the bottom chambers). To overcome the donor variability of switched IgE^+^ cells, we report the migration of cells in response to a specific chemokine as a percentage of cells that migrated in RPMI media (control).

### Analysis of chemokine receptor expression in nasal polyp scRNAseq

Raw scRNAseq reads from PCs isolated from NPs of patients with allergic fungal rhinosinusitis (National Institutes of Health SRA database BioProject number PRJNA898288) (23) were processed using CellRanger 6.0.1 (10x Genomics) and the GRCh38 genome reference. The resulting gene counts were analysed using Seurat 5.4.0 (28). Cells containing fewer than 200 or more than 6,000 detected genes, as well as cells containing more than 10% of mitochondrial gene transcripts, were excluded from analysis to remove low-quality cells, multiplets and dying cells, respectively. Data from each of the three donor samples were normalised using Seurat’s SCTransform function and integrated into a single dataset. The final analysis contained 9,244 cells. Genes encoding immunoglobulin variable regions and light chain constant regions (but not heavy chain constant regions) were excluded from the list of variable features before dimensionality reduction and clustering. The isotype of the immunoglobulin heavy chain expressed by each cell was identified using sciCSR (29). Briefly, the productively rearranged heavy chain isotype was determined as the isotype containing the most productive reads in each cell. Cells containing zero productive immunoglobulin heavy chain reads were classified as having isotype “Not Determined”. All plots were produced using the FeaturePlot and DotPlot functions in Seurat. Statistical analysis was performed using Seurat’s FindMarker function and grouping by isotypes.

## Results

### Human IgE^+^ PCs have reduced levels of CXCR4 and fail to migrate in response to CXCL12

To evaluate the chemokine receptor profile and the migration capacity of the human IgE-producing cells we class switched human tonsil B cells to IgE using IL-4 and anti-CD40. First, we determined the expression levels of CXCR4 in our switched IgE^+^ and IgG1^+^ cells on day 12 of the class switching cultures. As can be seen in Fig. 1 A, IgE-expressing cells have significantly lower levels of CXCR4 expression compared to IgG1^+^ cells. To understand the significance of the CXCR4 differential expression on IgE^+^ and IgG1^+^ cells, we tested their capacity to migrate in response to CXCL12 using the transwell chemotaxis assay system. To ensure absolute quantification of the migrating cells, we added the precision count beads to the bottom chamber of the transwell prior to harvesting and flow cytometry staining of the cells. Fig. S1 shows the gating strategy and the formula used to calculate the number of migrating cells. We accounted for the tonsil donor variability in the number of switched IgE^+^ and IgG1^+^ cells in cultures (26,27), by reporting the CXCL12 specific migration as a percentage of the cells migrating to the bottom chamber containing RPMI media control. Reflecting the levels of CXCR4 expression, we found that the migration of IgE^+^ cells in response to CXCL12 is significantly lower compared to that of all IgG1^+^ cells (Fig. 1 B).

**Figure 1.**
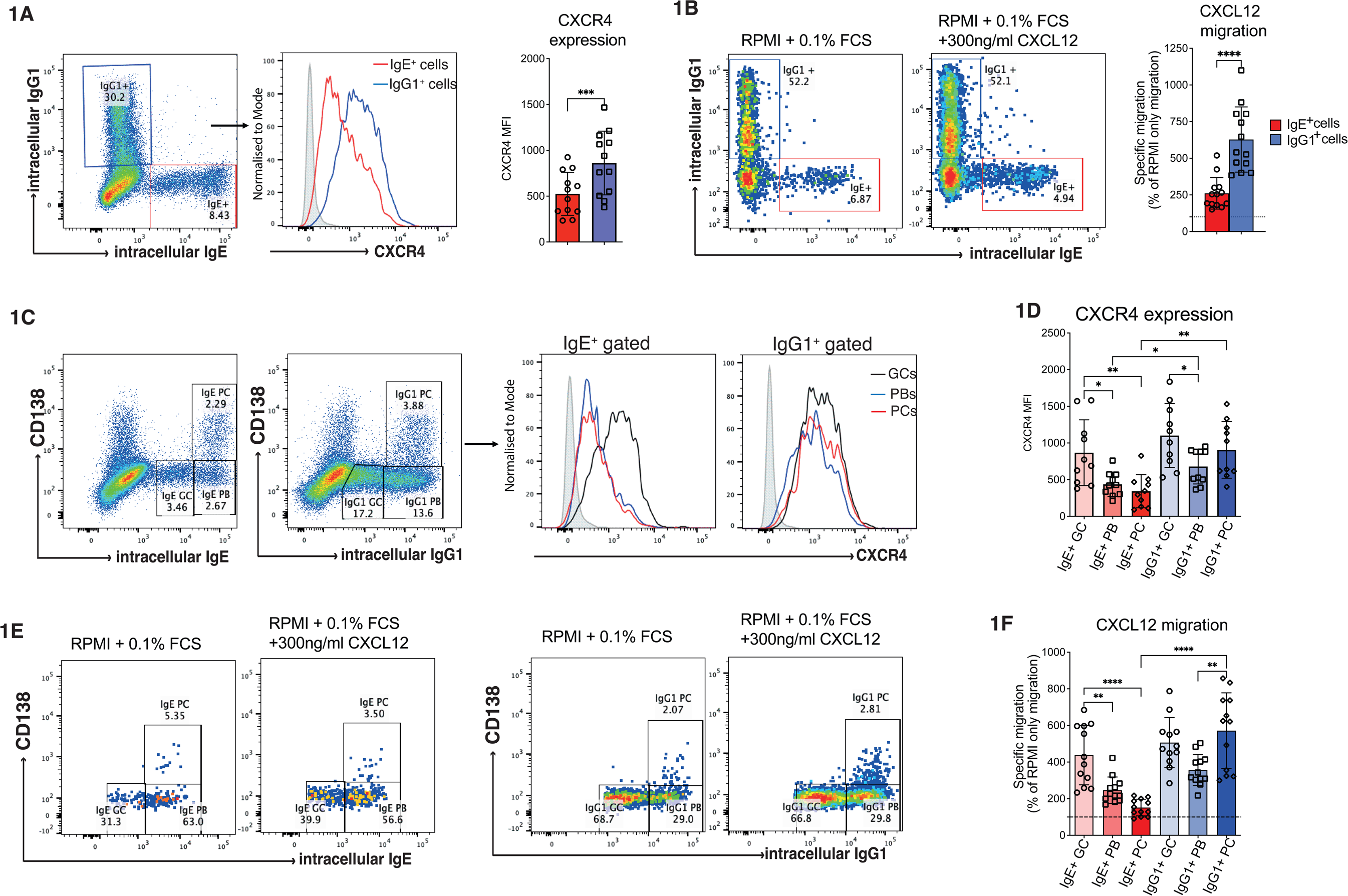
Human IgE^+^ PCs have reduced levels of CXCR4 and migrate poorly towards CXCL12. **(A)** Switched tonsil B cells were stained for surface CXCR4 followed by intracellular for IgE and IgG1. The histogram shows CXCR4 expression on IgE^+^ (red) and IgG1^+^ (blue) gated cells. The filled grey histogram represents the isotype control staining. The accompanying graph shows CXCR4 median fluorescence intensity (MFI) on the surface of IgE^+^ and IgG1^+^ cells. **(B)** CXCL12 induced migration of IgE^+^ and IgG1^+^ cells was assessed using the transwell assay. After 3h of migration, the number of migrating cells was quantified by flow cytometry. Migration of IgE^+^ and IgG1^+^ cells in response to 300 ng/mL (30nM) of CXCL12 is shown as a percentage of the cells migrating in response to RPMI control. **(C)** Flow cytometry dot plots of the IgE^+^ and IgG1^+^ gated GC-like B cells, a PC-like “plasmablast” and PCs. Representative histograms show the CXCR4 expression on each of these gated IgE^+^ and IgG1^+^ cells. **(D)** Bar chart showing the CXCR4 expression (MFI) across different IgE^+^ and IgG1^+^ cell populations. **(E)** Flow cytometry staining of IgE^+^ and IgG1^+^ GC-like B cells, PC-like PBs and PC after 3h of migration to the bottom chamber of the transwell. **(F)** CXCL12 induced migration shown as a percentage of the cells migrating in response to RPMI control. Each dot represents an individual tonsil donor. Data are mean + s.d. Statistical analysis was performed using one-way ANOVA with Tukey’s multiple comparison test (D, F) or paired two-tailed t-test with Welch’s correction (A, B); *p< 0.05; **p< 0.01; ***p< 0.001; and ****p< 0.0001. Non-significant values are not shown.

Previously, we reported three populations of IgE^+^ and IgG1^+^ cells in these *in vitro* class switching cultures; a GC-like B cell (IgE^lo^CD138^-^ and IgG1^lo^CD138^-^), a PC-like plasmablast (PB) (IgE^hi^CD138^-^ and IgG1^hi^CD138^-^) and a PC (IgE^hi^CD138^+^ and IgG1^hi^CD138^+^) population (27,30). Analysis of these populations shows that as IgG1^+^ cells differentiate from GC-like B cells to PBs and finally to PCs, they transiently downregulate the expression of CXCR4 at the PB stage before recovering it at the PC stage of differentiation (Fig. 1, C and D). In contrast, the differentiation of IgE^+^ GC-like B cells into PCs is followed by a significant down regulation of the CXCR4 expression (Fig. 1, C and D). In line with the CXCR4 expression on IgG1^+^ cells, we find that the IgG1^+^ GC-like B cells and IgG1^+^ PCs migrate to CXCL12 at similar rates, whereas the migration of IgG1^+^ PBs is reduced (Fig. 1, E and F). In addition, we also observe that the migration of IgE^+^ GC-like B cells to CXCL12 is comparable to that of IgG1^+^ GC-like B cells (Fig. 1, E and F). However, upon differentiation towards the PCs, IgE^+^ cells lose their migration response to CXCL12 (Fig. 1, E and F). These data confirm that IgE^+^ PCs migrate poorly in response to CXCL12 and affirm previous suggestions that CXCR4/CXCL12 recruitment of IgE^+^ PC into the BM may be impaired.

### IgE^+^ PCs selectively upregulate CCR2 and migrate in response to CCL2

Our transcriptional analysis of the IgE^+^ and IgG1^+^ cells previously identified two chemokine receptors that were significantly higher in IgE^+^ PCs compared to IgG1^+^ PCs (30). One of these receptors was CCR1, whose ligands, CCL3 and CCL5, are elevated in allergic rhinitis and during the allergic response in asthma individuals (31–33). However, when examining the surface CCR1 expression by flow cytometry we failed to detect any differences between IgE^+^ and IgG1^+^ cells along their differentiation into PCs (Fig. 2. A and B).

**Figure 2.**
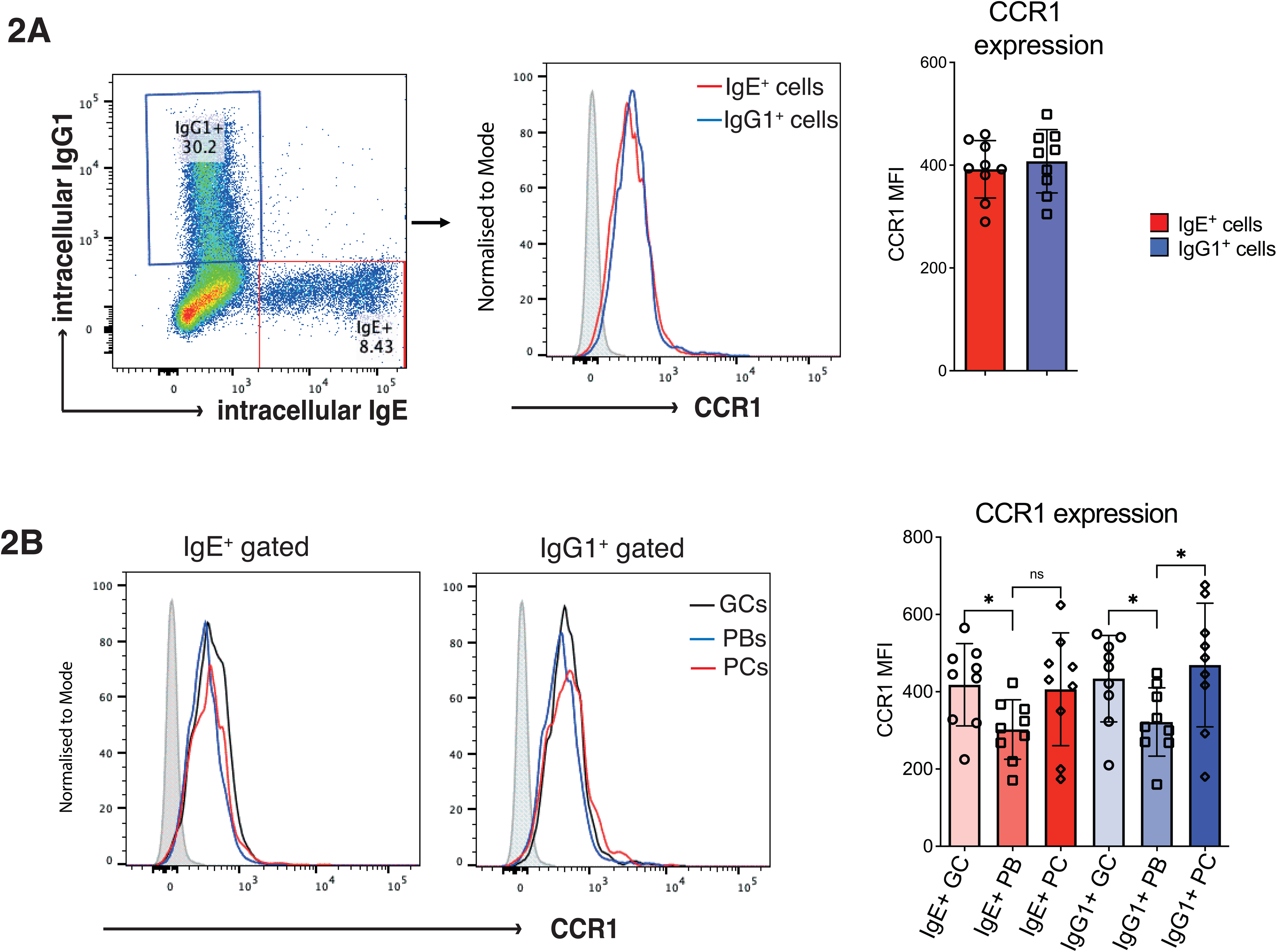
CCR1 expression on IgE^+^ and IgG1^+^ cells. **(A)** Representative histograms showing surface CCR1 expression on IgE^+^ gated cells (red) and IgG1^+^ gated cells (blue). The filled grey histogram represents the isotype control staining. The accompanying graph shows CCR1 MFI from the gated IgE^+^ and IgG1^+^ cells. **(B)** Histograms show the CCR1 expression on IgE^+^ and IgG1^+^ GC-like B cells, PBs, and PCs. CCR1 MFI across IgE^+^ and IgG1^+^ populations is shown in the column graph. Each dot represents an individual tonsil donor. Data are mean + s.d. Statistical analysis was performed using one-way ANOVA with Tukey’s multiple comparison test (B); *p< 0.05.

We identified CCR2 as the second chemokine receptor that was transcriptionally upregulated in human IgE^+^ PCs compared to IgG1^+^ PCs (30). CCL2, also known as the monocyte chemotactic protein 1 (MCP1), is the main ligand of CCR2, which is a potent activator and recruiter of monocytes and macrophages during inflammation (34,35). The CCL2/CCR2 axis also mediates the activation and recruitment of various other inflammatory cells playing a role in the pathogenesis of disease such as rheumatoid arthritis, multiple sclerosis, atherosclerosis and COVID-19(34,36,37). The CCL2/CCR2 axis can also be crucial to lung inflammatory responses in asthma (38–42). To determine whether CCR2 expression is also upregulated in IgE^+^ PCs derived from human tissues, we re-analysed a sc-RNAseq dataset from a previously published study of PCs from NPs of three patients with allergic fungal rhinosinusitis (23). Dimensionality reduction using uniform manifold approximation and projection (UMAP) identified 17 clusters of PBs and PCs in this dataset (Fig. 3 A). Determining the isotype of the productively re-arranged Ig heavy chain expressed in each cell revealed that IgE^+^ PCs resided primarily in cluster 7 (Fig. 3 B). Consistent with the *in vitro* transcription data, cluster 7 also had the highest expression of CCR2 (Fig. 3 C). Comparing CCR2 expression across Ig isotypes confirmed that IgE^+^ PCs have the highest CCR2 expression (*q*=0.00003 compared to PCs of all other isotypes, *q*=0.01637 compared to IgG1^+^ PCs) (Fig. 3 D). In contrast, CCR1 expression was mostly absent in the PCs from the NPs (Fig. S2 A, B, D).

**Figure 3.**
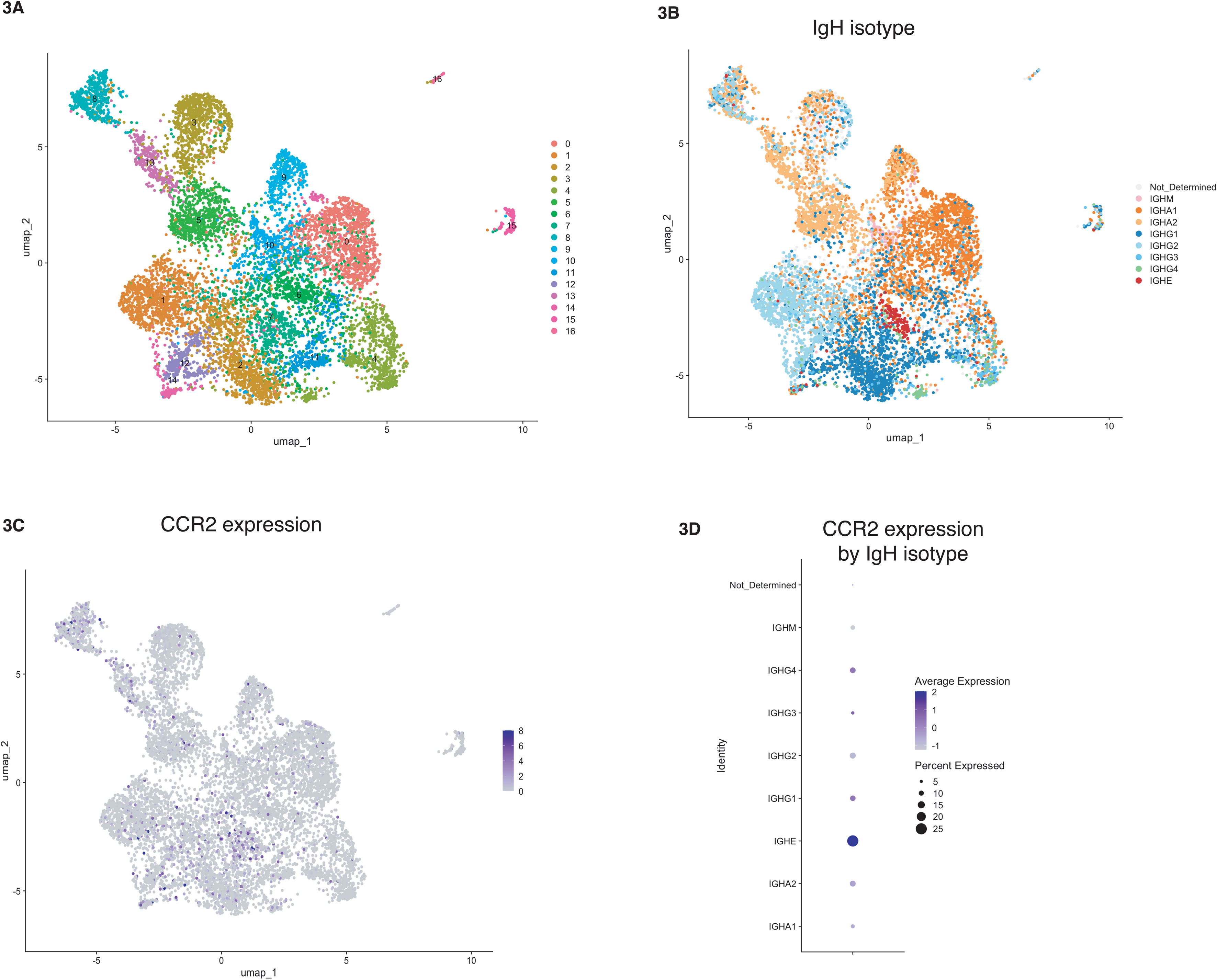
IgE^+^ PCs from NPs of patients with allergic fungal rhinosinusitis have significantly higher levels of CCR2 expression. **(A)** Two-dimensional uniform manifold approximation and projection (UMAP) plot illustrating the different clusters of PCs isolated from NPs of three patients with allergic fungal rhinosinusitis (23). **(B)** Expression of the productive IgH isotype in each cell overlaid onto the UMAP plot. IgE^+^ PCs (red) reside primarily in cluster 7 of the dataset. **(C)** Feature plot of log-normalized expression of CCR2 overlaid on the UMAP. expression. **(D)** Dot plot showing scaled CCR2 expression in PCs grouped by IgH isotypes.

We next confirmed that the CCR2 protein expression is significantly higher on the surface of IgE^+^ cells compared to IgG1^+^ cells by flow cytometry (Fig. 4 A). To test if IgE^+^ cells are more reactive to CCL2 compared with IgG1^+^ cells we measured their migratory capacity to three different concentrations of CCL2 (10 ng/mL, 100 ng/mL, and 300 ng/mL). As can be seen in Fig. 4 B, IgE^+^ cells migrate very potently even in response to 10 ng/mL (1.15 nM) of CCL2, and the IgE^+^ cell migration was further enhanced in response to 100 ng/mL (11.5 nM) and 300 ng/mL (34.5 nM) of CCL2. Reflecting their CCR2 expression, IgG1^+^ cells migrated very poorly in response to CCL2 (Fig. 4 B).

**Figure 4.**
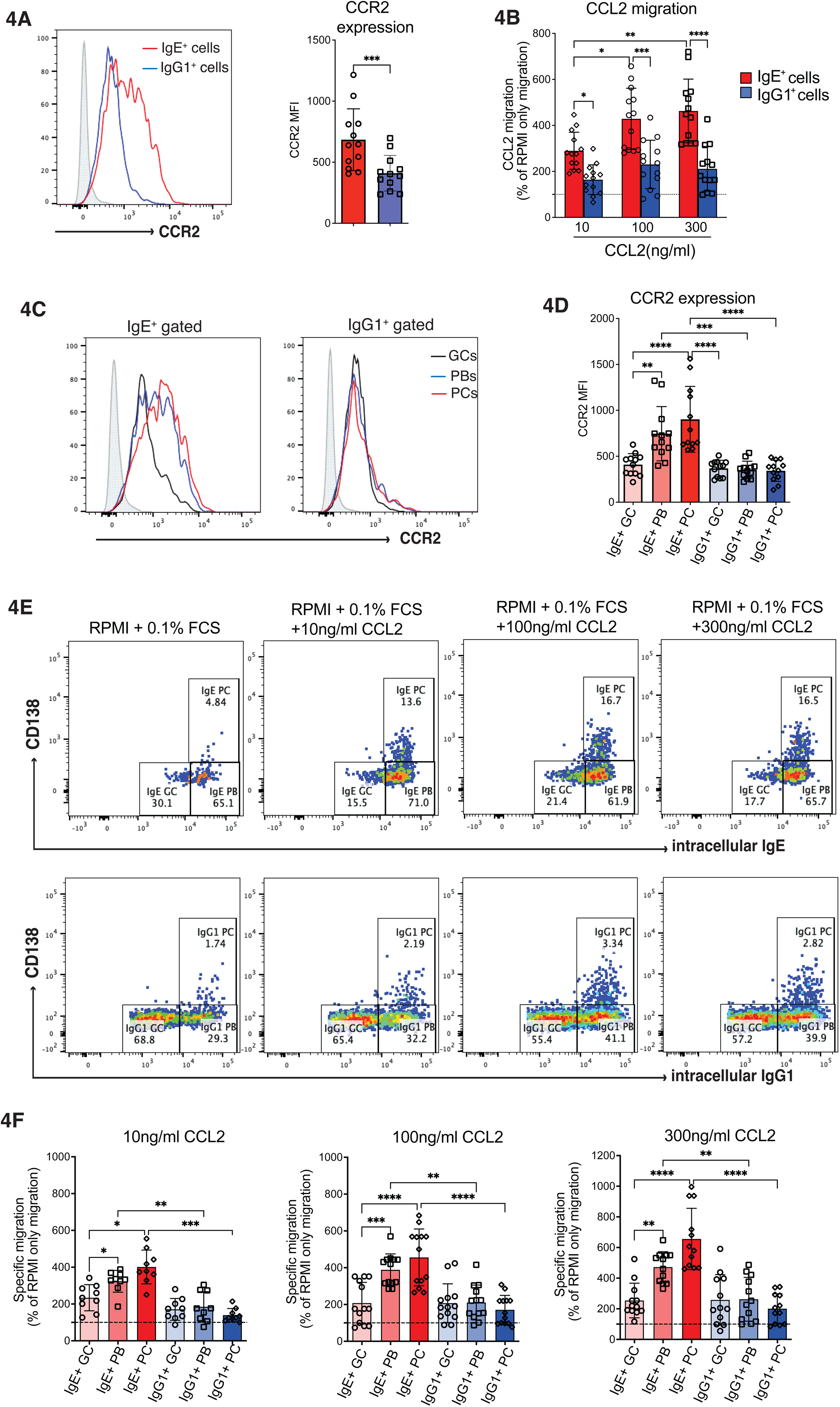
IgE^+^ PCs express high levels of CCR2 and migrate very efficiently in response to CCL2. **(A)** Representative histograms of CCR2 expression levels on IgE^+^ (red) and IgG1^+^ (blue) gated cells. The filled grey histogram represents the isotype control staining. The accompanying graph shows CCR2 MFI on IgE^+^ and IgG1^+^ gated cells. **(B)** Migration of IgE^+^ and IgG1^+^ cells in response to 10 ng/mL (1.15nM), 100 ng/ml (11.5nM) and 300 ng/mL (34.5nM) of CCL2, shown as a percentage of the cells migrating in response to RPMI control. **(C)** Histograms of CCR2 expression on IgE^+^ and IgG1^+^ gated GC-like B cells, a PC-like “plasmablast” and PCs. **(D)** CCR2 MFI across the different IgE^+^ and IgG1^+^ cell populations from several experiments. **(E)** Dot plot visualisation of IgE^+^ and IgG1^+^ GC-like B cells, PB and PC after 3h of migration to the bottom chamber of the transwell containing different concentrations of CCL2. **(F)** CCL2 induced migration of the different IgE^+^ and IgG1^+^ cell populations shown as a percentage of the cells migrating in response to RPMI control. Each dot represents an individual tonsil donor. Data are mean + s.d. Statistical analysis was performed using one-way ANOVA with Tukey’s multiple comparison test (B, D, F) or paired two-tailed t-test with Welch’s correction (A); *p< 0.05; **p< 0.01; ***p< 0.001; and ****p< 0.0001.

When examining the CCR2 expression in IgE^+^ and IgG1^+^ cells at different stages of their PC differentiation, we observed a significant increase in the levels of CCR2 expression in both IgE^+^ PBs and IgE^+^ PCs compared with IgE^+^ GC- like B cells (Fig. 4, C and D). In contrast, IgG1^+^ cell populations maintain similar levels of CCR2 expression along their differentiation pathway, suggesting that CCR2 is selectively upregulated in IgE^+^ PCs. Reflecting these differential levels of CCR2 expression, we find that both IgE^+^ PBs and IgE^+^ PCs migrate towards CCL2 at a significantly higher rate compared to IgE^+^ GC-like B cells (Fig. 4, E and F). As expected, IgG1^+^ cells migrated very poorly in response to CCL2 (Fig. 4, E and F). Taken together, these data demonstrate that while the migration of human IgG1^+^ PCs is predominately dependent on CXCL12 and CXCR4 interaction, CCR2 and CCL2 interaction is a key regulator of the IgE^+^ PB and IgE^+^ PC migration, suggesting that IgE^+^ PCs have a distinct homing preference from IgG1^+^ PCs.

### Both IgE^+^ and IgG1^+^ cells upregulate CCR10 upon PC differentiation and migrate at similar rates in response to CCL28

CCL28, also known as mucosae-associated epithelial chemokine (MEC), is constitutively expressed by epithelial cells in various mucosal tissues, including the gastrointestinal, respiratory and salivary glands (43–45). Through its binding to its receptor, CCR10, CCL28 has also been implicated in the regulation of IgE^+^ cell migration (46). Our previous work showed that CCR10 is elevated in IgE^+^ PCs when compared with IgE^+^ GC-like B cells at the transcription level (30), suggesting that IgE^+^ PCs might indeed migrate more readily in response to CCL28. However, the transcription data also showed that there were no differences between IgE^+^ and IgG1^+^ cells. This is supported by the flow cytometry data in Fig. 5 A and B, which show an increase in the surface CCR10 expression on both IgE^+^ and IgG1^+^ PCs compared to GC-like B cells and PBs, but no differences in CCR10 expression between IgE^+^ and IgG1^+^ expressing cells. The sc-RNAseq analysis of PCs from NPs showed that CCR10 is predominantly expressed by IgA2^+^, IgA1^+^ and IgM^+^ PCs in this tissue (Fig. S2 A, C and D). The IgG1^+^ and IgE^+^ PCs from the NPs showed similarly modest expression of CCR10 (Fig. S2 A, C and D). Next, we tested the migration capacity of IgE^+^ and IgG1^+^ cells in response to CCL28. We find that both IgE^+^ and IgG1^+^ cells failed to migrate in response to 300 ng/mL (25 nM) of CCL28, which is the same as the optimal concentrations of CCL2 and CXCL12 used (Fig. 5, C-F). However, when using a higher concentration of CCL28, 1.5 μg/mL (125 nM), we observed a significant increase in the migration of both IgE^+^ and IgG1^+^ cells (Fig. 5, C and D). Further analysis confirmed that IgE^+^ PCs and IgG1^+^ PCs, in line with their increased CCR10 expression, migrated towards CCL28 at a significantly higher rate than their less differentiated cell predecessors (Fig. 5, E and F). This data suggests that some IgE^+^ and IgG1^+^ PCs, which express higher levels of CCR10, may home towards CCL28 in tissues.

**Figure 5.**
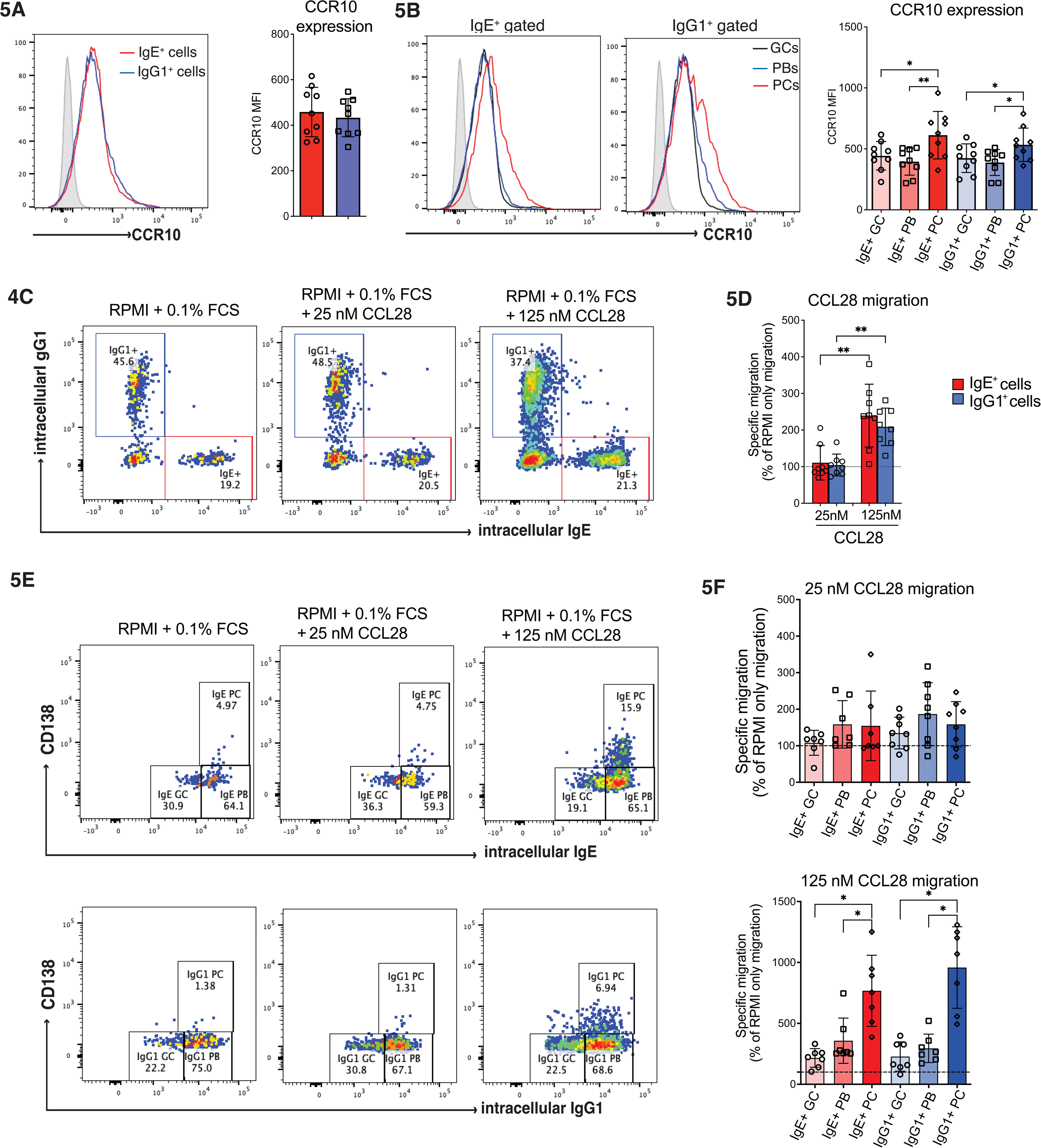
IgE^+^ and IgG1^+^ PCs upregulate CCR10 and migrate at similar rates in response to CCL28. **(A)** Representative histograms showing surface CCR10 expression on IgE^+^ (red) and IgG1^+^ (blue) gated cells; the filled grey histogram shows the isotype control staining. The accompanying graph shows CCR10 MFI on the gated IgE^+^ and IgG1^+^ cells. **(B)** Representative plots of CCR10 expression on IgE^+^ and IgG1^+^ GC-like B cells, PBs, and PCs. CCR10 MFI across the different IgE^+^ and IgG1^+^ cell populations from several experiments. **(C)** The dot plots show the frequency of IgE^+^ and IgG1^+^ cells after 3h of migration in response to RPMI only or two different concentrations of CCL28, 25nM (300 ng/mL) and 125nM (1.5 μg/mL). **(D)** CCL28 induced migration shown as a percentage of the IgE^+^ and IgG1^+^ cells migrating in response to RPMI control. **(E)** The migration capacity of IgE^+^ and IgG1^+^ GC-like B cells, plamablasts, and PC in response to CCL28. **(F)** CCL28 induced migration shown as a percentage of the cells migrating in response to RPMI control. Each dot represents an individual tonsil donor. Data are mean + s.d. Statistical analysis was performed using one-way ANOVA with Tukey’s multiple comparison test (B, D, F) or paired two-tailed t-test with Welch’s correction (A); *p< 0.05; **p< 0.01.

## Discussion

Migration of plasma cells to survival niches in the BM and other tissues is crucial for the maintenance of long-lived antibody responses. In the case of IgE^+^ PCs, their lifespan has been the focus of considerable discussion, yet little is known about the migratory and homing preferences that underpin their persistence, particularly in humans. Early observations that mouse IgE^+^ PCs have impaired migration in response to CXCL12 (6), a chemokine that regulates cell migration into the BM niches (16,17), led to the suggestion that these cells fail to populate the BM survival niches and are therefore limited in their longevity.

Consistent with these observations, we find that human IgE^+^ PCs derived from our culture migrate poorly towards CXCL12 compared to IgG1_⁺_ PCs. Mechanistically, the loss of this migration correlates with significantly lower CXCR4 surface expression on these cells, suggesting that the reduced CXCR4 expression on IgE^+^ PCs may selectively restrict their access to the BM niches, thereby limiting the accumulation of these cells within this compartment. It is important to note, however, that these findings may not fully recapitulate the diversity of CXCR4 expression states reported across in vivo settings (6,7,18,19,23). These discrepancies are likely to reflect tissue and context specific differences in CXCR4 regulation, potentially shaped by the local environment in which IgE_⁺_ PCs differentiate and reside. However, in line with their impaired migration in response to CXCL12, the IgE^+^ PC numbers present in the BM are very low (7,18–20).

Based on our data, which show that the less differentiated IgE_⁺_ cells in our cultures migrate towards CXCL12 at levels comparable to their IgG1_⁺_ counterparts, it is tempting to speculate that early IgE_⁺_ PC precursors retain some capacity for BM recruitment before downregulating CXCR4. Once established within the BM niches, IgE^+^ PCs may subsequently recover CXCR4 expression that helps promote their long-term retention and survival. Consistent with this, IgE^+^ PCs detected in the BM of mice after 15 weeks of chronic allergen inhalation initially express reduced surface CXCR4 relative to IgG1^+^ PCs but recover CXCR4 expression to levels like those of IgG1^+^ PCs after a further 9 weeks following cessation of allergen exposure (7). Given the established role of CXCR4-CXCL12 interactions in PC retention and survival, such recovery of CXCR4 expression may help stabilise these allergen-specific IgE^+^ PCs in BM survival niches and support their continuous secretion of allergen-specific IgE antibodies into the circulation. In line with this, the few IgE^+^ PCs that do establish in the BM following chronic allergen exposure are long-lived since they are resistant to cyclophosphamide treatment (5) and persist after B cell depletion (7).

The most novel finding of this study is the selective upregulation of CCR2 on IgE_⁺_ PCs and their robust migration towards its ligand, CCL2. In contrast, IgG1^+^ PCs fail to upregulate CCR2 and migrate poorly in response to CCL2, highlighting an Ig isotype specific migration programme. This selectivity was observed at both the protein level by flow cytometry and at the transcriptional level by re-analysis of a published scRNAseq dataset from human NPs (23), where IgE_⁺_ PCs showed the highest CCR2 expression among all PC isotypes.

The scRNAseq study of IgE_⁺_ PCs from NPs of allergic donors also provided validation of our in vitro observations in a physiologically relevant human tissue. Considering that CCR2 regulates migration of cells to inflamed tissues (34), it is plausible that migration in response to CCL2 may enable IgE^+^ PCs to preferentially home to sites of inflammation during allergen exposure. Indeed, several reports have shown that allergen challenges elevate the levels of CCL2 in the bronchoalveolar lavage (BAL) fluid of asthma individuals (38,41,42) and of animal models of allergic asthma (41,47,48). Blocking CCL2 with a neutralising antibody also helps reduce IgE production in BAL during ovalbumin induced lung allergic inflammation (49), suggesting that CCL2 interaction with CCR2 may also help regulate IgE responses during lung inflammatory responses. Importantly, in addition to migration, CCL2 binding to CCR2 can also activate signalling pathways involved in cell survival (50), raising the possibility that homing of IgE^+^ PCs to CCL2-rich tissues not only determines their localisation but may also support their persistence. While this remains to be determined, such a mechanism would provide an explanation for the maintenance of IgE^+^ PCs outside classical BM niches.

We also considered whether CCR2 signalling might influence CXCR4-mediated migration and BM retention of IgE_⁺_ PCs. The precedent for such CCR2-CXCR4 crosstalk comes from earlier studies showing that CCR2 activation on immature B cells reduces the CXCR4 mediated migration (51). In addition, CCR2 activation can also desensitise CXCR4 signalling on cells already in the BM, which disrupts the anchoring provided by the CXCL12-CXCR4 interaction, leading to their mobilisation into the blood stream. This has been shown in BM monocytes during the inflammatory response (52). In fact, CCR2 signalling is known to regulate the egress of various other leukocytes from the BM (53). Even in the absence of an inflammatory response, inhibition of CXCR4 leads to the exit of the CCR2 expressing cells from the BM (52). Similarly, blocking CXCR4 has been reported to enhance the egress of PCs from the BM (54). Thus, by analogy, activation of CCR2 on IgE_⁺_ PCs may both recruit these cells to the inflammatory tissues and simultaneously dampen their CXCR4-dependent anchoring within BM niches. Due to the reduced CXCR4 in our IgE^+^ PCs, the CCR2-CXCR4 crosstalk in IgE_⁺_ PCs has not been tested experimentally in this study, and its relevance to IgE_⁺_ PC biology remains to be determined

Another important finding we report here is the upregulation of CCR10 by a fraction of IgE^+^ PCs, and migration of these cells towards CCL28, which has been reported to be involved in allergic airway inflammation and asthma (55–57). However, in contrast to the CCR2, we did not detect any differences between IgE^+^ and IgG1^+^ PCs with respect to CCR10 expression or their migratory response to CCL28. In addition to CCL28, which is expressed by the epithelial cells of various mucosal tissues (43), CCR10 has another ligand, CCL27, which is predominantly expressed in the skin (58) where it plays a crucial role in inflammatory skin diseases like atopic dermatitis (59,60). There is a significant correlation between the severity of atopic dermatitis and higher levels of IgE, which is increased in both serum and skin of patients (61,62). While some of the IgE-producing cells might be generated locally, our data suggest that CCR10 interaction with its ligands, CCL27 and CCL28, may also facilitate the homing of the CCR10^+^IgE^+^ PCs to the skin and various other mucosal tissues.

Taken together, our findings highlight a previously unknown key regulator of IgE^+^ PC migration and provide insights that help explain both the limited accumulation of IgE_⁺_ PCs in the BM and their preferential localisation to peripheral tissues. We discovered that, instead of CXCR4, IgE^+^ PCs specifically express high levels of CCR2, which facilitates their migration towards its ligand, CCL2. This key discovery lays the foundation for future in vivo studies to determine the functional importance of CCR2 in regulating localisation and longevity of IgE^+^ PCs in human allergic disease.

## Acknowledgments

We are grateful to the patients and the ENT surgical team at the Guy’s & St Thomas’ NHS Foundation Trust for their help and support in the collection of tonsils used in this research. We also thank Dr David Fear (King’s College London) for their support with the collection of tonsils used in this research. We thank Dr Astrid Fabri and Dr Adam McShane (Institute of Immunity and Transplantation, UCL) for helpful comments on the manuscript. This study was supported by Asthma UK career development award (AUK-CDA-2019-412) to F.R and Wellcome Trust Investigator Award (223196/Z/21/Z) to P.T.

## Author Contributions

Z.L. investigation, analysis, methodology, and writing—review and editing. P.T. investigation, analysis, methodology, funding acquisition, and writing—review and editing. F.R. conceptualization, investigation, analysis, methodology, funding acquisition, and writing—original draft, review, and editing.

## Conflict-of-interest

The authors declare that they have no conflicts of interests.

### Abbreviations

BM: Bone Marrow
BAL: Bronchoalveolar lavage
CCR1: Chemokine (C-C motif) receptor 1
CCR2: Chemokine (C-C motif) receptor 2
CCR10: Chemokine (C-C motif) receptor 10
CXCR4: Chemokine (C-X-C motif) receptor 4
CCL2: Chemokine (C-C motif) ligand 2
CCL28: Chemokine (C-C motif) ligand 28
CXCL12: Chemokine (C-X-C motif) ligand 4
GC: Germinal Centre
PB: Plasmablast
PC: Plasma cells
UMAP: Uniform Manifold approximation and projection

