## Supplementary figure 1 and 2 for "IgE-producing cells on the move: CCR2 is a key regulator of IgE^+^ plasma cell migration"

### Supplementary Figures

## S1A

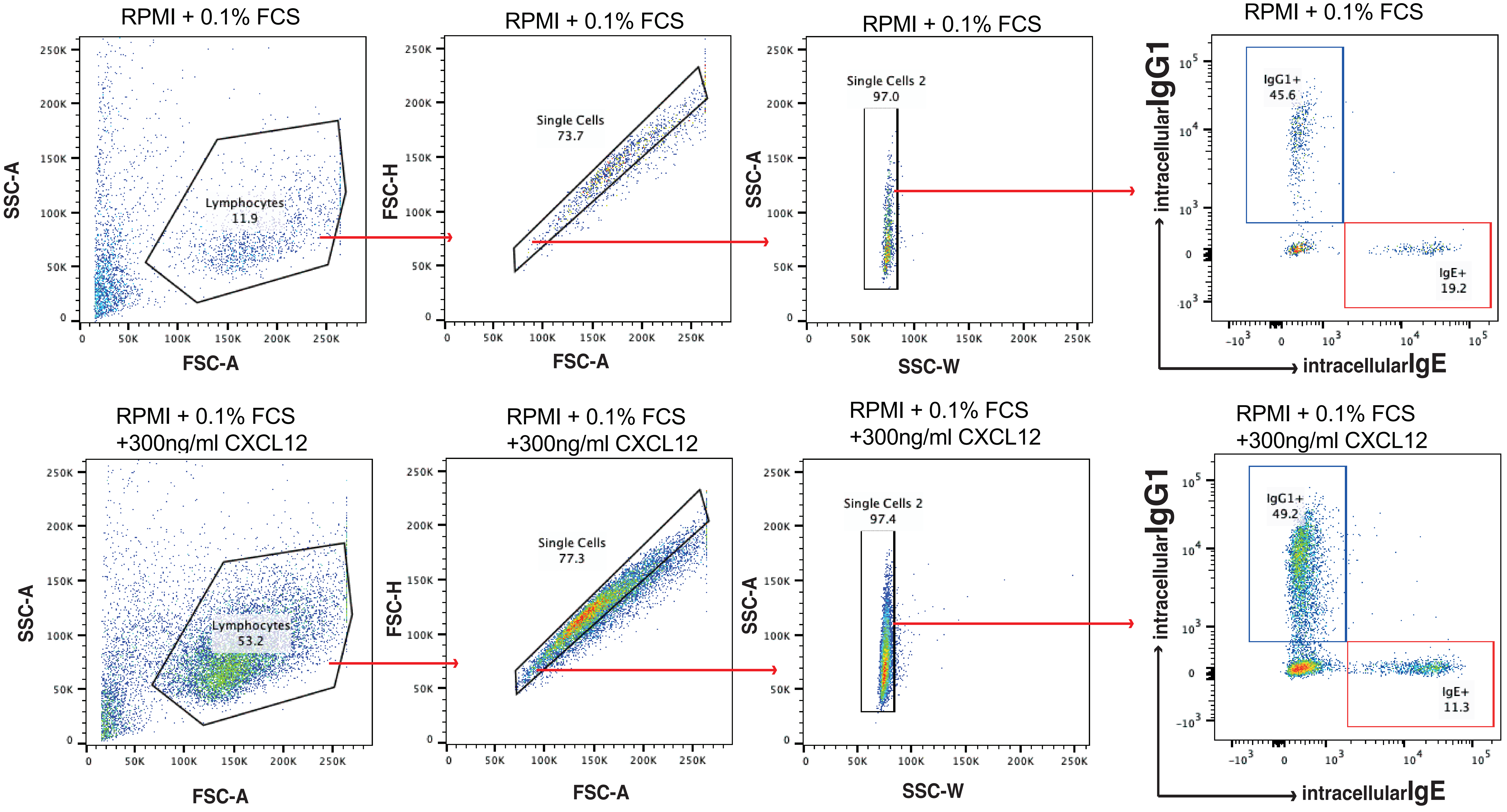


## S1B

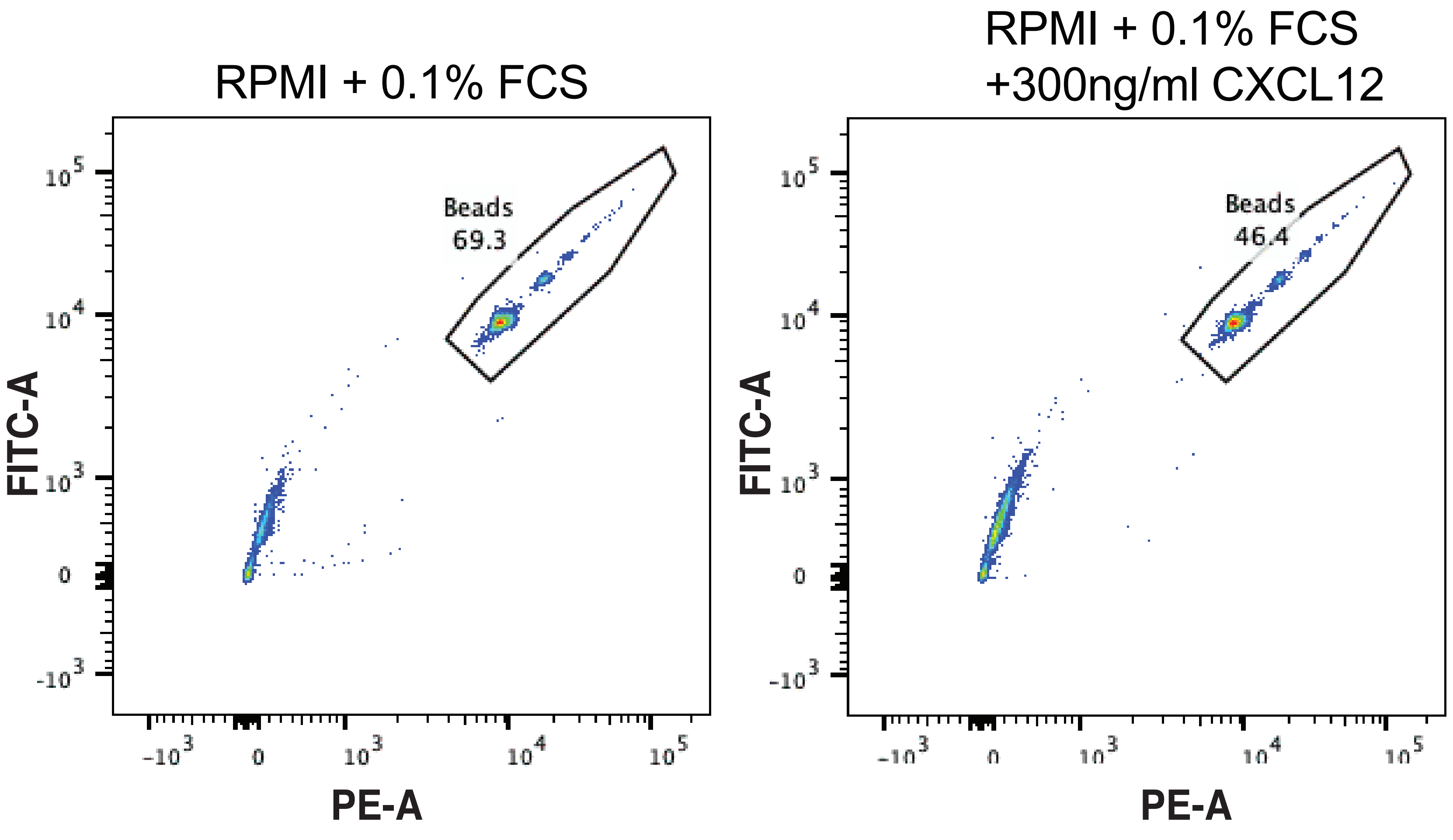


## S1C

$$\text{Number of migrated cells} = \frac{\text{Number of IgE}^{+}\text{ or IgG1}^{+}\text{ cells in the bottom well}}{\text{Number of counted beads}} \times \text{Total number of beads added}$$

**Figure S1. Overview of the flow cytometry and analysis workflow.**

**(A)** The gating strategy used in our analysis of $IgE^+$ and $IgG1^+$ cells.

**(B)** To ensure absolute quantification of the migrated cells we also gated the precision counting beads, which were added to the bottom chamber before cell harvesting and staining.

**(C)** Formula used to calculate the number of migrated cells recovered from the bottom chamber.

S2A

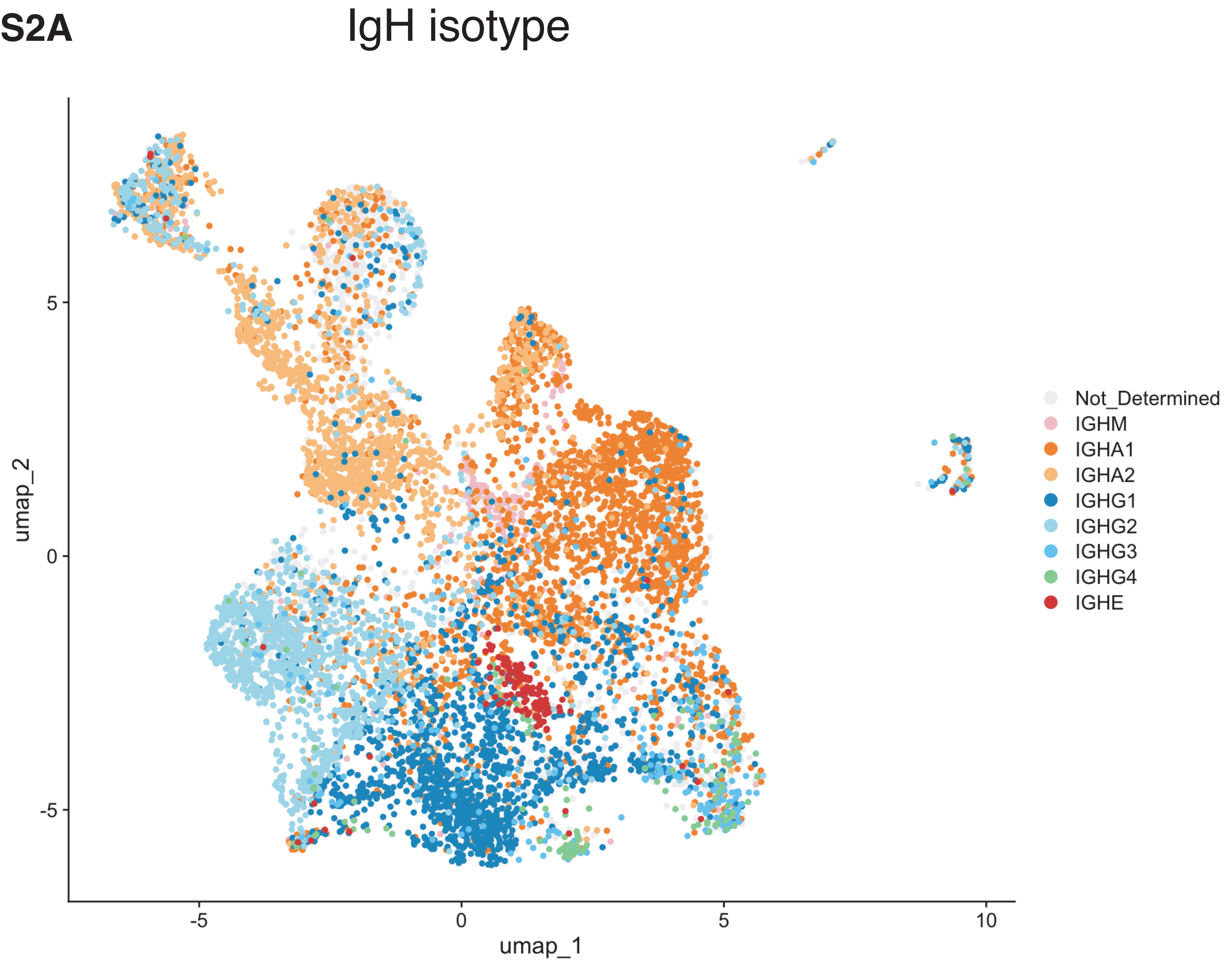


S2B

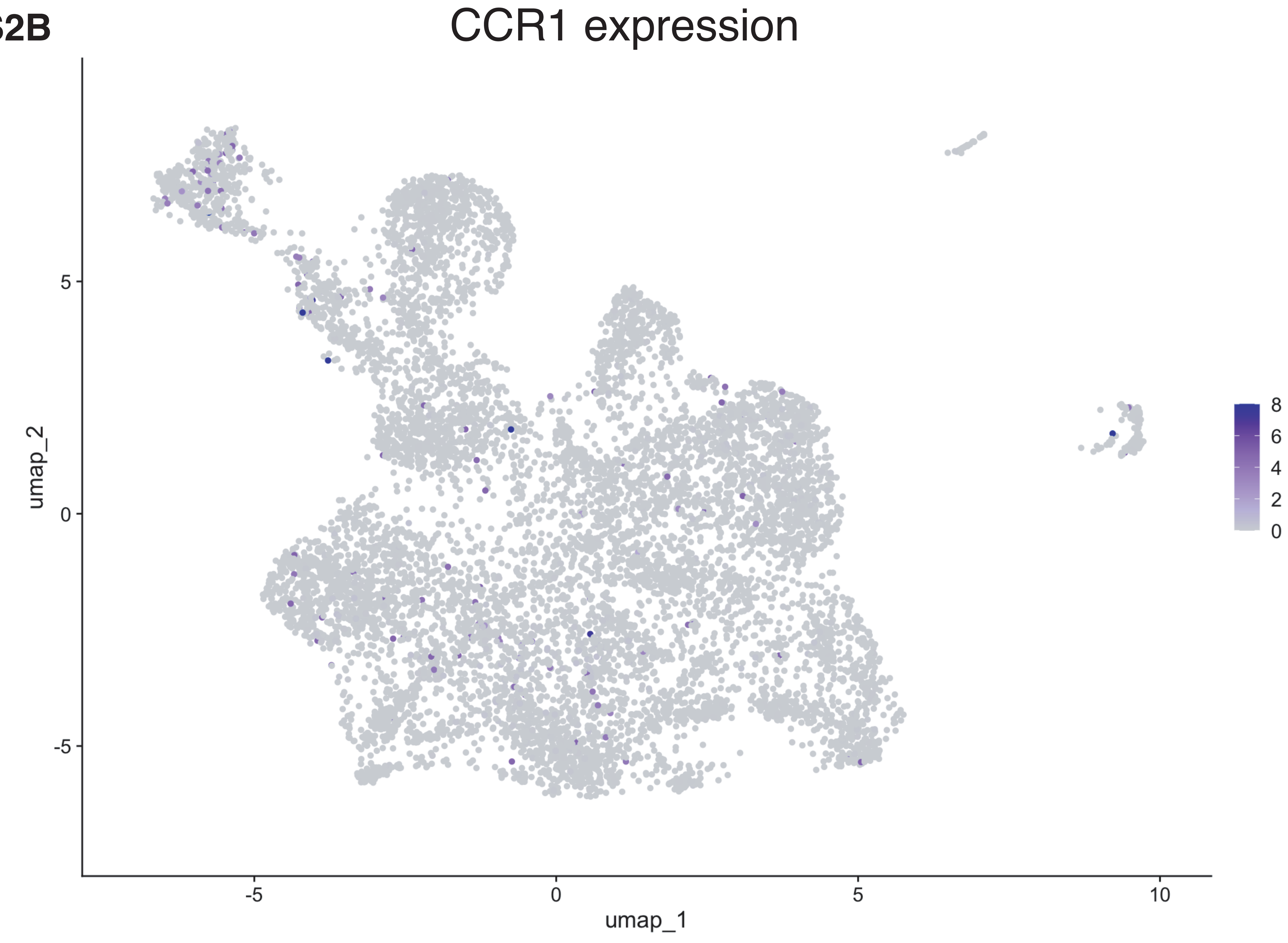


S2C

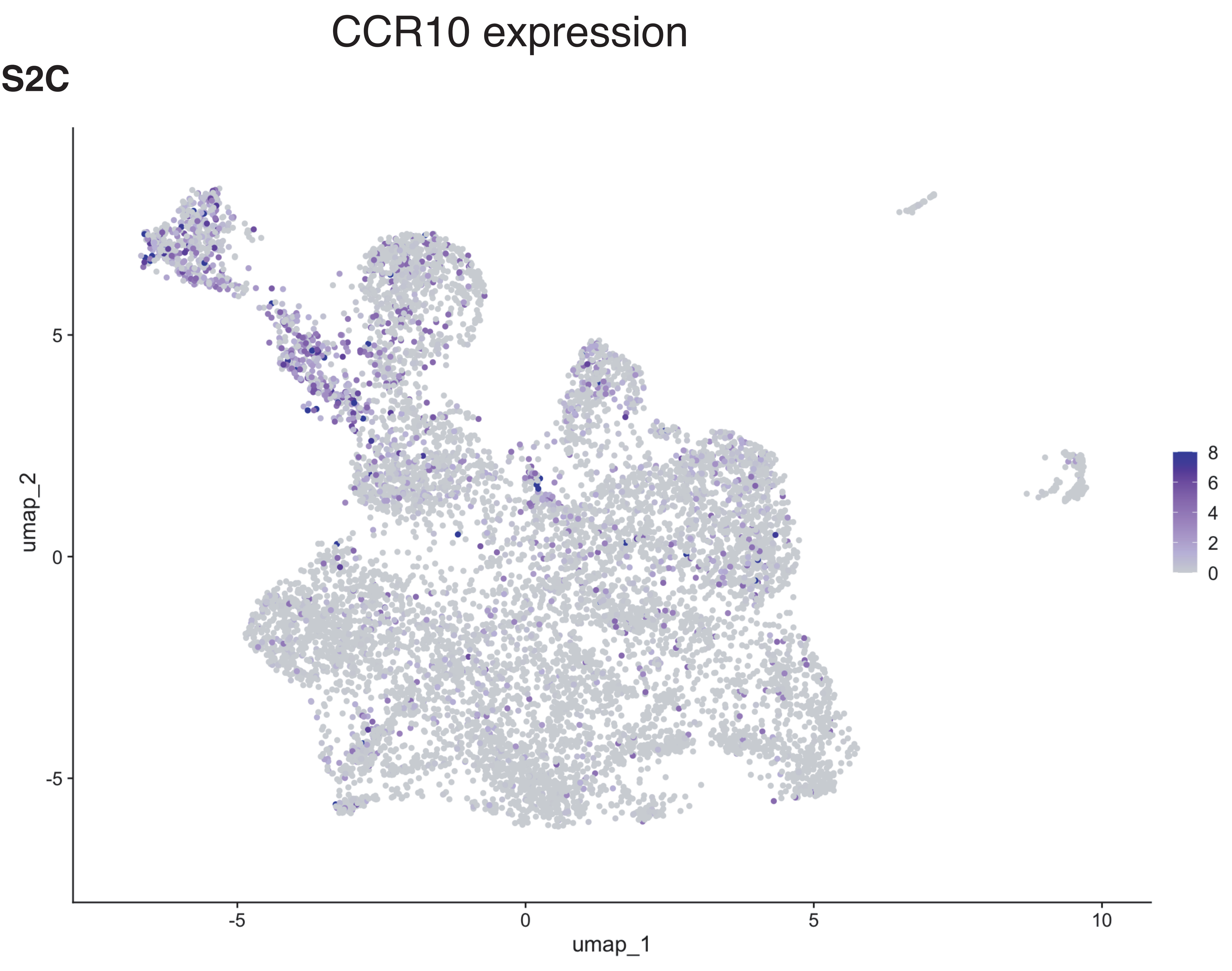


S2D

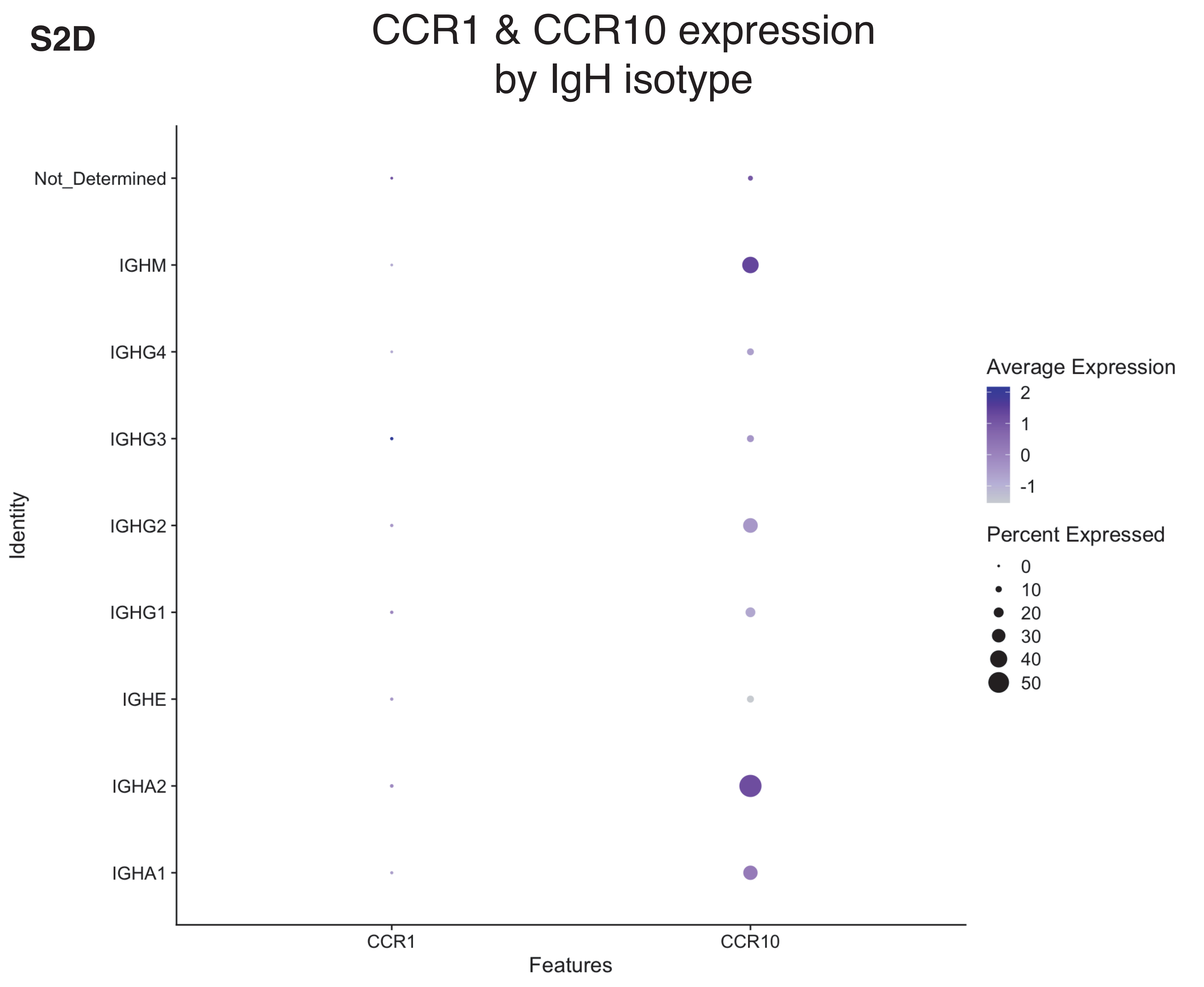

**Figure S2. CCR1 and CCR10 expression in PCs from nasal polyps of patients with allergic fungal rhinosinusitis.**

**(A)** Expression of the productive IgH isotype in each cell overlaid onto the UMAP plot.

**(B)** Feature plot of log-normalized expression of CCR1 overlaid on the UMAP. expression.

**(C)** Feature plot of log-normalized expression of CCR10 overlaid on the UMAP. expression.

**(D)** Dot plot showing scaled CCR1 and CCR10 expression in PCs grouped by IgH isotypes.
